# UBL3 interaction with α-synuclein is downregulated by silencing MGST3

**DOI:** 10.1101/2023.05.15.540904

**Authors:** Jing Yan, Hengsen Zhang, Bin Chen, Yashuang Ping, Md. Shoriful Islam, Yuna Tomochika, Shuhei Aramaki, Tomohito Sato, Yu Nagashima, Tomohiko Nakamura, Tomoaki Kahyo, Daita Kaneda, Kenji Ogawa, Minoru Yoshida, Mitsutoshi Setou

## Abstract

Ubiquitin-like 3 (UBL3) is a membrane-anchored protein that has been discovered to function as a protein post-translational modifier, helping to sort proteins into small extracellular vesicles (sEVs). Aggregations of alpha-synuclein (α-syn) are associated with the pathology of neurodegenerative diseases such as Parkinson’s disease (PD). The aggregation and toxicity of α-syn can be influenced by its interactions with specific proteins. Recently, the interactions between UBL3 and α-syn was uncovered. It is believed to play a role in eliminating excess α-syn from neurons, or in the spread of α-syn pathology in the brain associated with neurodegenerative diseases. However, the regulator that can mediate the interaction between UBL3 and α-syn remains unclear. In this study, we employed the split gaussian luciferase complementation assay and RNA interference (RNAi) technology to discover that QSOX2, HTATIP2, UBE3C, MGST3, NSF, HECTD1, SAE1 and ATG3 are involved in downregulating the interaction between UBL3 and α-syn. Among these proteins, silencing MGST3 had the most significant impact on the UBL3-α-syn interaction (with a fold change log2 of less than - 1). MGST3 is a part of the antioxidant system, and silencing MGST3 is believed to contribute to oxidative stress. We used hydrogen peroxide (H_2_O_2_) to induce oxidative stress and observed its effect on the UBL3-α-syn interaction. Our finding showed that an 800μM concentration of H_2_O_2_ could also downregulate this interaction. However, the effect of oxidative stress caused by 800μM H_2_O_2_ on the UBL3-α-syn interaction did not be enhanced by silencing MGST3. In conclusion, the interaction between UBL3 and α-syn is downregulated by silencing MGST3.

## Introduction

Ubiquitin-like 3 (UBL3), also known as membrane-anchored ubiquitin-fold protein (MUB), is a highly conserved ubiquitin-like protein [1]. In 2018, it was discovered that UBL3 can serve as a post-translational modifier of proteins, tagging them for sorting into small extracellular vesicles (sEVs) [2]. sEVs are secreted by various cells with a diameter of 30-150nm [3]. They play crucial roles in mediating cell-to-cell communication in both physiology and pathology. For instance, sEVs that transport proteins between cells have been implicated in the development of neurodegenerative diseases [4].

Alpha-synuclein (α-syn) is encoded by the *SNCA* gene consisting of 140 amino acids and weighing 14.5kDa. It is highly expressed in the adult central nervous system especially in presynaptic terminals [5]. Abnormal aggregations of α-syn are a major feature of Lewy bodies which are found in neurons of patients with neurodegenerative diseases such as sporadic Parkinson’s disease (PD) and Dementia with Lewy bodies (DLB) [6,7]. Research has shown that the interactions of α-syn with specific proteins can influence its aggregation and toxicity. For example, β-amyloid peptides can interact with α-syn and promote the intraneuronal accumulation of α-syn [8] whereas heat shock protein 70 (Hsp70) can interact with α-syn and aid in its clearance thereby reducing its aggregation [9]. Modulation of these interactors is emerging as a promising therapeutic strategy for neurodegenerative diseases.

Recently, the interaction of UBL3 with α-syn has been revealed using split gaussian luciferase complementation assay and immunoprecipitation [10]. This interaction is thought to be related to the clearance of excess α-syn from neurons, or the propagation of α-syn pathology in the brain associated with neurodegenerative diseases. However, the regulator involved in regulating the interaction of UBL3 with α-syn are currently unknown.

In this study, we used split gaussian luciferase complementation assay and RNA interference (RNAi) technology to discover silencing MGST3 downregulate the interaction of UBL3 with α-syn.

## Materials and Methods

### Plasmid and siRNA

The NGluc-UBL3 and α-syn-CGluc carried by the pCI vector, and Gluc plasmid were used in the previous study of our laboratory [10]. The siRNAs used for silencing target candidate proteins were purchased from Silencer Select (Ambion, Life Technologies, CA, USA) (Supplementary Table 1). Two siRNAs were available for each candidate protein. One additional siRNA was provided as a negative control group to assure silencing efficiency and avoid siRNA toxic effect.

### Cell culture and Transfection

Human embryonic kidney (HEK293) cells (RIKEN Cell Bank, Japan) were cultured in Dulbecco’s Modified Eagle Medium (DMEM, GIBCO, 11965-092) supplemented with 10% fetal bovine serum (FBS) at 37°C in a humidified incubator with 5% CO_2_.

The cells were seeded in 24-well plate. When the cell confluency reached 70-80%, transfection was performed using lipofectamine 2000 (Thermo Fisher Scientific, Waltham, MA, USA) and Opti-MEM reduced serum medium (Thermo Fisher Scientific, Waltham, MA, USA). Specifically, 100ng of NGluc-UBL3 plasmid, 100ng of α-syn-CGluc plasmid, and 20 pmol of siRNA were transfected per well. The transfection was performed according to the manufacturer’s instruction for lipofectamine 2000.

### Luciferase assay

Cells were incubated for 72 hours after transfection. We collected the cell culture medium and centrifuged it at 1200 rpm for 5 min to remove cell debris. Then, the supernatant was added to 17 μg/mL coelenterazine (Cosmo Bio, Kyodo, Japan) diluted by Opti-MEM medium and luminescence was immediately measured using BioTek Synergy H1 microplate reader (Agilent).

### Western blot

Cells were washed with ice-cold PBS and harvested by centrifugation at 2000 × g for 5 min at 4 °C. Then, the cells were resuspended and lysed with 1% Triton lysate buffer (50 mM Tris-HCl [pH 7.4], 100 mM NaCl, and 1% [v/v] Triton X-100) for 30 min on ice. Cell lysate was centrifuged at 20,000 × g for 15 min at 4 °C to remove cell debris and un-lysed cells. Quantification of protein concentration was performed using the Pierce BCA Protein Assay Kit (23227, Thermo Fisher Scientific, IL, USA). For WB analysis, 10 μg of total protein was loaded after being treated with 2-mercaptoethanol (βME) sodium dodecyl sulfate (SDS) sample loading buffer (100 mM Tris-HCl [pH 6.8], 4% SDS, 20% glycerol, and 0.01% bromophenol blue) at 95°C for 5 min, separated on SDS-PAGE gel and transferred to polyvinylidene fluoride (PVDF) membrane. The membrane was blocked with 5% skim milk for 1 hour at room temperature and then were incubated overnight at 4°C with the anti-MGST3 antibody (Abcam, ab192254; 1: 1000). Subsequently it was incubated with horseradish peroxidase (HRP)-conjugated secondary antibody (Cell signaling, 1:5000 dilution) for 1 hour at room temperature after washing. We detected the protein signal using the Enhanced Chemiluminescence Kit (32106, Thermo Fisher Scientific, Waltham, MA, USA) with the FUSION FX imaging system (Vilber Lourmat, Collégien, Seine-et-Marne, France).

### Oxidative stress

We used hydrogen peroxide (H_2_O_2_) (FUJIFILM, Japan) to induce oxidative stress. After transfection, the cells were passaged to 96-well plate with 100 μL of DMEM (10% FBS) per well and incubated overnight. The H_2_O_2_ was diluted with pre-warmed DMEM (10% FBS). We replaced 50 μL of culture medium in each well with the culture medium containing different concentrations of H_2_O_2_. The final H_2_O_2_ concentrations were 0μM, 100μM, 200μM, 400μM and 800μM. Cells were further incubated 48 hours, and then the culture medium was collected for luciferase assay.

### MTT Assay

The 3-(4,5-dimethylthiazol-2-yl)-2,5-diphenyl-2H-tetrazolium Bromide (MTT) cell growth kit (CT02, Millipore, MA, USA) was used for measuring cell viability. After collecting cell culture medium for luciferase assay, 100μL pre-warmed DMEM (10% FBS) medium was added to each well with 10 μL of MTT reagent to incubate 4 hours. Then we added 100 μL solubilization buffer, isopropanol with 0.04 N hydrogen chloride, and the absorbance value (OD) at 450nm wavelength was measured using the microplate reader. Cell viability equal to (OD Intervention group – OD Blank group) / (OD Control group – OD Blank group). The blank group had only medium without cells and the control group had medium and cells without intervention.

### Statistical analysis

Statistical analysis was performed using GraphPad Prism 8.0 software. The results from three independent experiments are presented as the mean ± S.D. Statistical significance was assessed by two-tailed Student’s t-test for two groups. *p*<0.05 was considered statistically significant.

## Results

### Screening of regulators affecting the interaction of UBL3 with α-syn using split gaussian luciferase complementation assay and RNAi technology

In previous study to discover that UBL3 modification affects the sorting of proteins to the sEVs, they performed proteomic analysis to identify proteins that interact with UBL3 in a manner dependent on two c-terminal cysteine residues [2]. From these proteins, we selected 10 candidate proteins based on modification functions involving ubiquitination, glycosylation, acetylation, etc. that may affect the interaction of UBL3 with α-syn. We purchased corresponding siRNAs to silence these 10 candidate proteins (Supplement Table 1) and co-transfected them with NGluc-UBL3 and α-syn-CGluc into HEK293 cells respectively. After 72 hours of incubation, we collected the cell culture medium to detect the luminescence (Figure 1A). Compared to the control group transfected with only NGluc-UBL3 and α-syn-CGluc, the luminescence was significantly downregulated in the group transfected with siRNA silencing QSOX2, HTATIP2, UBE3C, MGST3, NSF, HECTD1, SAE1 and ATG3. After being transformed by fold-change log2 (Figure 1B), we found silencing MGST3 has the greatest effect on this interaction (Fold change Log2<-1). Thus, the QSOX2, HTATIP2, UBE3C, MGST3, NSF, HECTD1, SAE1 and ATG3 are involved in regulating the interaction of UBL3 with α-syn, where the MGST3 has a greatest effect.

**Figure 1.**
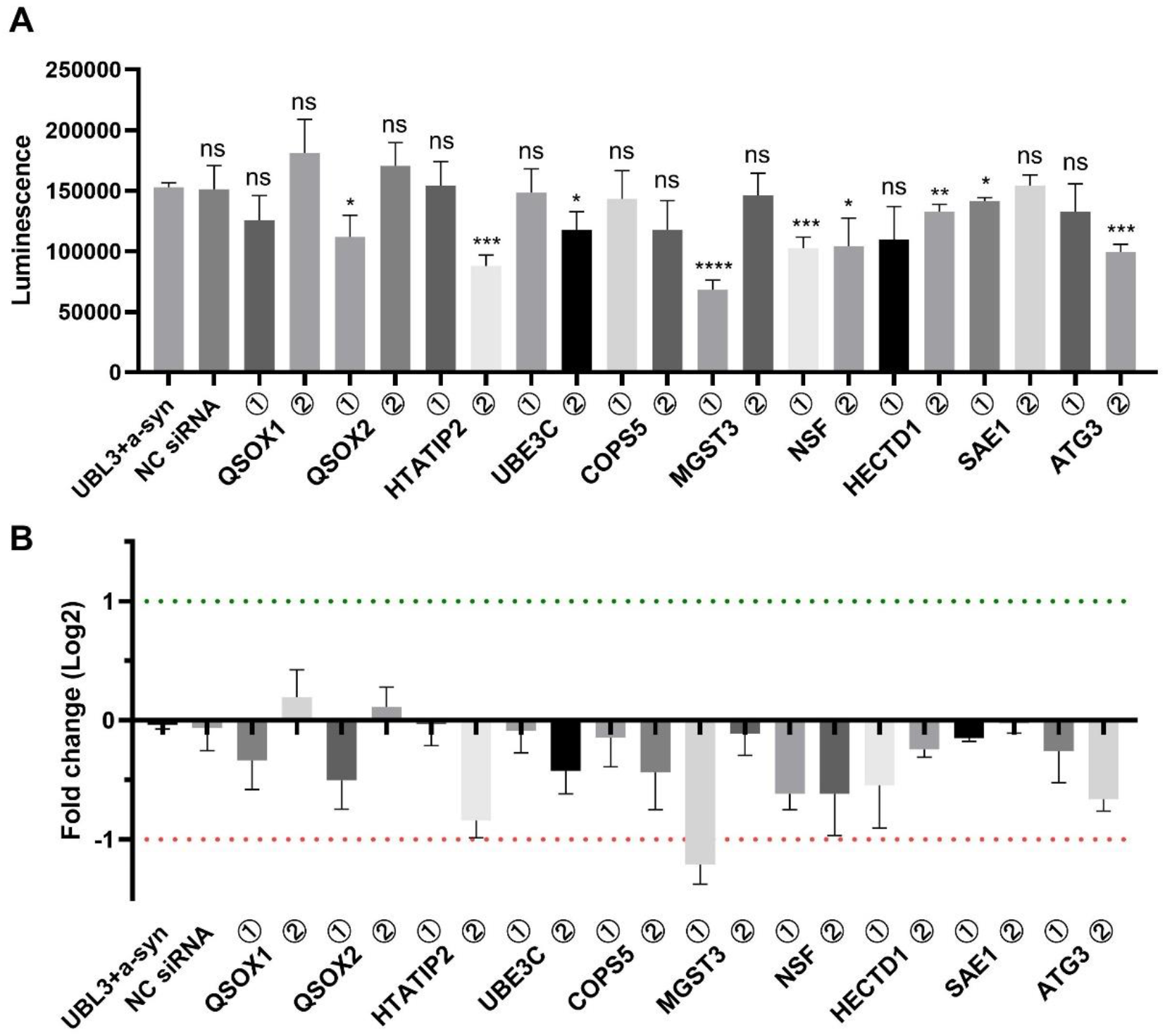
Effect of siRNA on the interaction of UBL3 with α-syn using split-luciferase complementation assay. (A) luminescence of culture medium of HEK293 cells co-transfected with siRNA, NGluc-UBL3 and α-syn-CGluc. Two siRNAs were available for each candidate protein. The groups were compared with the UBL3 and α-syn co-transfected group, and statistically analyzed. (B) Fold change log2 of luminescence. Silencing MGST3 resulted in luminescence fold change log2<-1. The luminescence ± S.D. in three independent experiments as shown. ns: non-significant; *: *p*<0.05; **: *p*<0.01, ***: *p*<0.001, ****: *p*<0.0001. NC siRNA: negative control siRNA, QSOX1: quiescin sulfhydryl oxidase 1, QSOX2: quiescin sulfhydryl oxidase 2, HTATIP2: HIV-1 Tat interactive protein 2, UBE3C: ubiquitin protein ligase E3C, COPS5: COP9 signalosome subunit 5, MGST3: microsomal glutathione S-transferase 3, NSF: N-ethylmaleimide sensitive factor, HECTD1: HECT domain E3 ubiquitin protein ligase 1, SAE1: SUMO1 activating enzyme subunit 1, ATG3: autophagy related 3

### Confirmation of MGST3 expression silencing

After quantification of the protein from the cell lysate, we performed western blot analysis to confirm the knockdown of MGST3 expression (Figure 2). We compared the signal of MGST3 in the group transfected with MGST3 siRNA to the control group and found that the signal was attenuated in the MGST3 siRNA transfected group. It indicated successful knockdown of MGST3 by siRNA in the cells.

**Figure 2.**
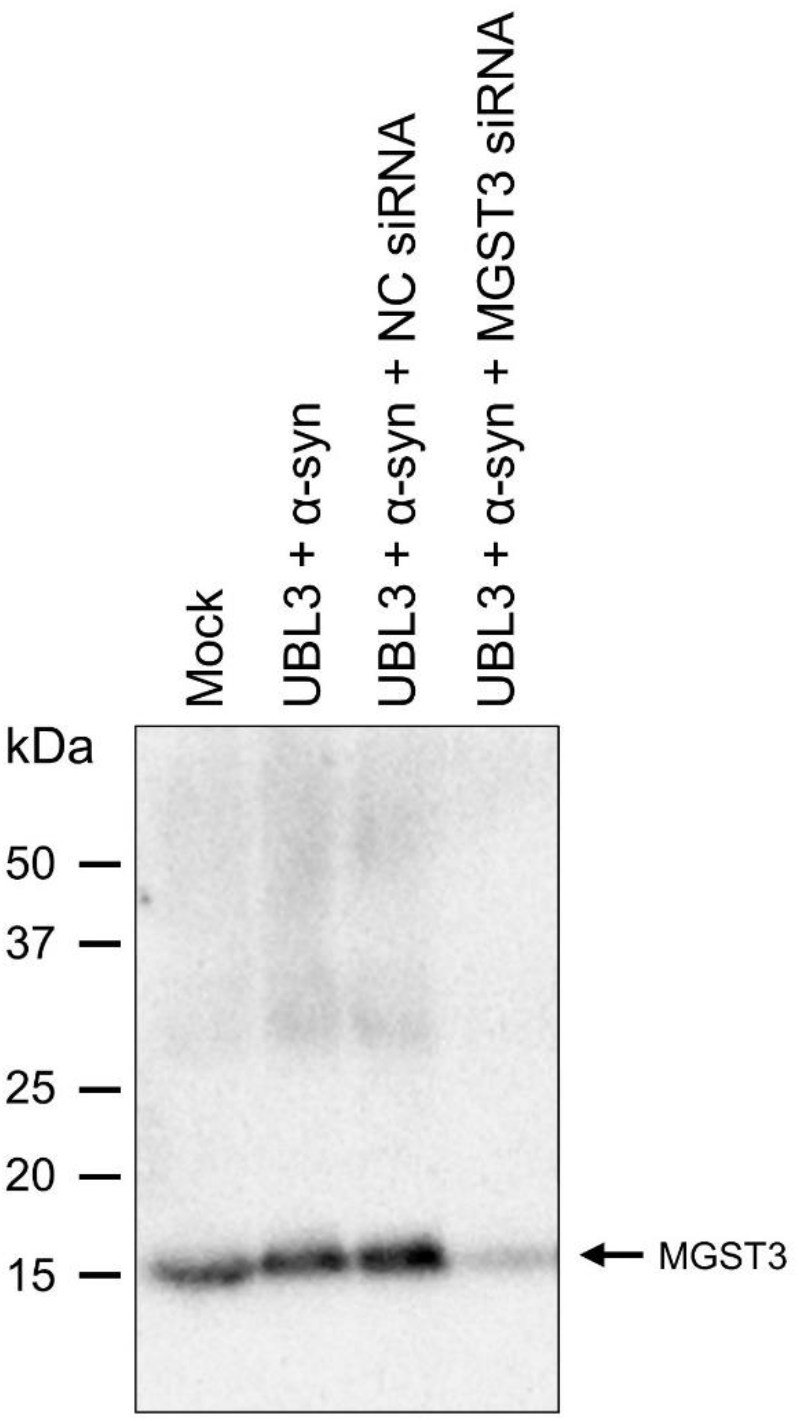
The MGST3 was knockdown of HEK293 cells transfected with MGST3 siRNA. NC siRNA: negative control siRNA.

### Effect of oxidative stress on the interaction between UBL3 and α-syn

MGST3 has mainly glutathione transferase and glutathione peroxidase activities [11,12]. It is well-known to play a protective role against oxidative stress [13]. Therefore, we verified whether oxidative stress affects the interaction of UBL3 with α-syn.

The oxidative stress in vitro model has been proposed by using H_2_O_2_ as an inducer when it is added to the cell culture medium [14]. We used cell culture medium containing different concentrations of H_2_O_2_ to culture HEK293 cells transfected with NGluc-UBL3 with α-syn-CGluc. Afterward, cell culture medium was collected and the luminescence was measured (Figure 3A). Only the 800μM H_2_O_2_ treated group showed a significant decrease in luminescence compared to the 0μM H_2_O_2_ treated group. Meanwhile, we observed that the H_2_O_2_ exhibited potent cytotoxicity at a concentration of 800μM, leading to partial cell death. Therefore, we used MTT assay to detect cell viability (Figure 3B). Consistent with our observation, the result of MTT assay showed a significant decrease in cell viability when H_2_O_2_ reached 800μM. To reduce the effect of cytotoxicity of H_2_O_2_ on the result, we calculated the ratio of luminescence to the cell viability. When the H_2_O_2_ concentration was 800μM, the ratio decreased significantly compared to the control, while other lower concentrations had no effect (Figure 3C). We treated cells transfected with Gluc using 800μM H_2_O_2_ and we found that the ratio of luminescence to the cell viability were not different from the group without treating with 800μM H_2_O_2_ (Figure 3D). The effect of H_2_O_2_ on luciferase activity was excluded. This suggested that a certain level of oxidative stress can downregulate the interaction between UBL3 and α-syn.

**Figure 3.**
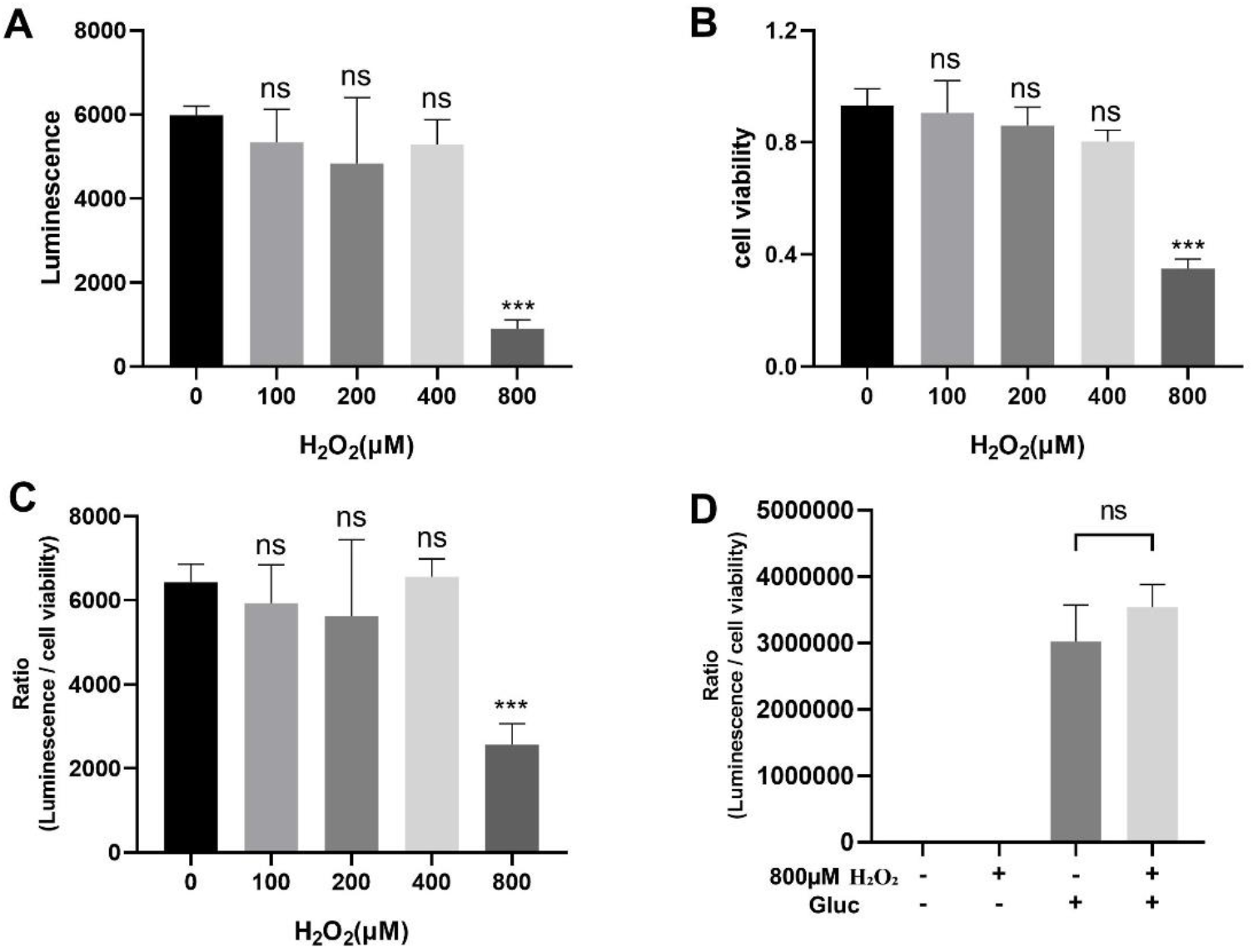
Effect of H_2_O_2_ on the interaction between UBL3 and α-syn. (A) luminescence of culture medium from HEK293 cells co-transfected with NGluc-UBL3 and α-syn-CGluc at different concentrations of H_2_O_2_. (B) Effect of different concentrations of H_2_O_2_ on the viability of HEK293 cells co-transfected with NGluc-UBL3 and α-syn-CGluc. (C) The ratio of luminescence to cell viability of culture medium from HEK293 cells co-transfected with NGluc-UBL3 and α-syn-CGluc at different concentrations of H_2_O_2_. These groups were compared to the group treated with 0μM H_2_O_2_, and statistically analyzed. (D) The ratio of luminescence to cell viability of culture medium from HEK293 cells transfected with Gluc treated with 800μM H_2_O_2_. The luminescence ±S.D., cell viability ±S.D. and ratio ±S.D. in three independent experiments as shown. ns: non-significant, ***: *p* < 0.001.

### Effect of silencing MGST3 on the interaction of UBL3 with α-syn upon oxidative stress

Silencing MGST3 could contribute to oxidative stress in cells. It remains unclear whether silencing MGST3 can exacerbate the effects of oxidative stress on UBL3-α-syn interactions or not. To verify this, we treated with H_2_O_2_ at a concentration of 800μM after performing co-transfection of NGluc-UBL3, α-syn-CGluc, and MGST3 siRNA. Cell culture medium was obtained after 48 hours of incubation and assayed for luminescence (Figure 4A). MTT assay was also used to determine the cell viability and the ratio of luminescence to cell viability was calculated (Figure 4B, C). By statistical analysis, we observed that after silencing MGST3, the interaction of UBL3 with α-syn was further downregulated after treatment with H_2_O_2_ at a concentration of 800μM, as shown in Figure 4C. However, there was no difference of the ratio of luminescence to cell viability between the groups treated with H_2_O_2_ in the presence or absence of silent MGST3. This indicated that silencing MGST3 did not affect the downregulation of interaction between UBL3 and α-syn by oxidative stress.

**Figure 4.**
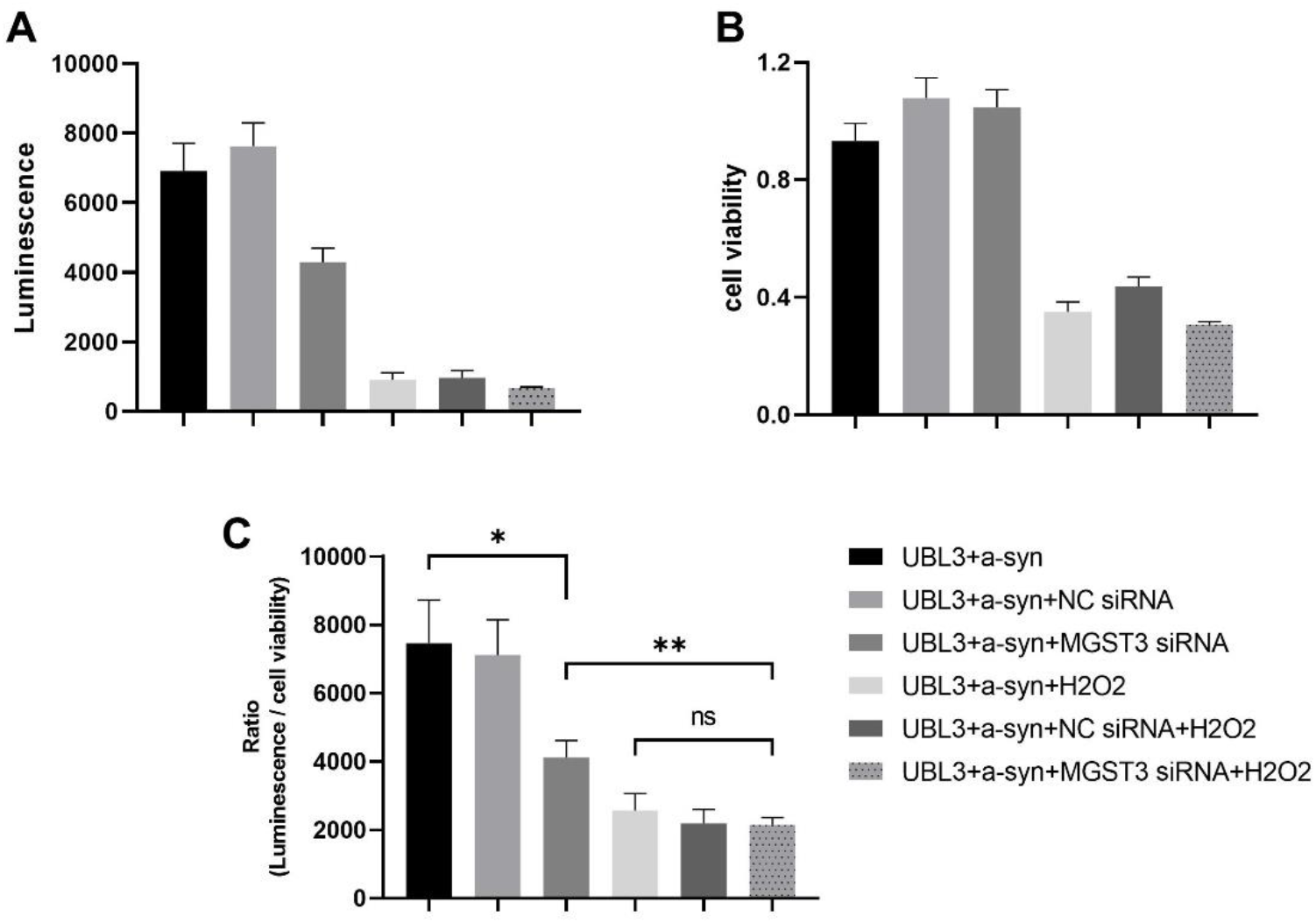
Effect of silencing MGST3 on the interaction between UBL3 and α-syn with 800μM H_2_O_2_. (A) luminescence of culture medium from HEK293 cells co-transfected NGluc-UBL3 with α-syn-CGluc. (B) Viability of the HEK 293 cells. (C) The ratio of luminescence to cell viability of HEK293 cell culture medium co-transfected with NGluc-UBL3 and α-syn-CGluc. The luminescence ±S.D., cell viability ±S.D. and ratio ±S.D. in three independent experiments as shown. NC siRNA: negative control siRNA, ns: non-significant, * *p* < 0.05, ** *p* < 0.01.

## Discussion

In this study, we found that silencing QSOX2, HTATIP2, UBE3C, MGST3, NSF, HECTD1, SAE1 and ATG3 significantly downregulated the interaction of UBL3 with α-syn where silencing MGST3 have the greatest effect among them.

As an enzyme belonging to the MAPEG (membrane associated proteins in eicosanoid and glutathione metabolism) family [15], MGST3 catalyzes the conjugation of glutathione to electrophilic compounds, facilitating their removal from cells, and participates in reduction of H_2_O_2_ and lipid peroxides to their corresponding alcohols [11]. This activity helps to protect cells from oxidative damage and maintain redox homeostasis. It is also commonly used as a biomarker for oxidative stress [16,17]. MGST3 is widely expressed in various human tissues, including the heart, brain, liver, kidney, pancreas, thyroid, skeletal muscle, testis, and ovary [11]. A study used data from a genome-wide association study (GWAS) of brain structure in humans identified a significant positive correlation between MGST3 expression and the size of the hippocampus [18], and the authors suggested that dysregulation of MGST3 may be associated with neurodegenerative diseases. The aggregation of α-syn is one of the typical pathological features of neurodegenerative diseases, and its pathogenesis involves multiple factors, including neuroinflammation, oxidative stress, protein metabolism disorders, etc. [19,20]. Although MGST3 plays a certain role in these factors, there is currently no study that clearly demonstrates that MGST3 affects the aggregation of α-syn. Our finding provides a promising starting point for further investigation of the potential link between MGST3 and α-syn aggregation.

We conducted a screening of proteins that interact with UBL3 and identified MGST3 as a protein that affects the interaction of UBL3 with α-syn. Our results showed that H_2_O_2_ can downregulate the interaction of UBL3 with α-syn. Based on this, we hypothesized that MGST3 can form a complex, MGST3-UBL3-α-syn. The presence of MGST3 can use glutathione to break down H_2_O_2_ in the vicinity of the complex, thereby protecting the UBL3-α-syn interaction from being downregulated by H_2_O_2_. When MGST3 is silenced, cellular oxidative imbalance and increased H_2_O_2_ generation, resulting in downregulation of the interaction of UBL3 with α-syn. Interestingly, under the condition of H_2_O_2_ treatment, we did not observe a further reduction in the interaction of UBL3 with α-syn by MGST3 silencing. We speculate that this may be due to the fact that the intensity of H_2_O_2_ stimulation in this experiment is beyond the range that endogenous MGST3 can effectively regulate. In the future, it would be interesting to investigate whether overexpression of MGST3 can prevent the downregulation of the UBL3-α-syn interaction by H_2_O_2_.

It is still unclear how H_2_O_2_ alters the interaction of UBL3 with α-syn based on the information currently available. The C-terminal cysteine residues of UBL3 are essential for the post-translational modification of the target protein with UBL3 [2], but that modification seemed not to occur to the interaction of UBL3 with α-syn [21]. It is possible that H_2_O_2_ may affect the intracellular interaction of UBL3 with α-syn by altering the covalent modification of UBL3 with other unknown factors mediated by the C-terminal cysteine residues.

Oxidative stress caused by the excessive production of reactive oxygen species (ROS) such as H_2_O_2_, hydroxyl radicals and superoxide from cellular metabolic processes leads to cell damage. Growing evidences implicate that the oxidative stress is one of the crucial factors involved in the pathogenesis of neurodegenerative diseases [22,23]. For example, the brain of PD patients exhibits decreased mitochondrial function and excessive production of ROS, including depletion of endogenous antioxidants and oxidative damage to cellular macromolecules such as proteins, lipids, and nucleic acids [24,25]. It has been found that the production of ROS can trigger the formation of Lewy bodies [26]. In addition, increased levels of cholesterol metabolites of ROS have been reported in the cerebral cortex of patients with DLB, and it has also been shown that increased levels of such metabolites accelerate the aggregation of α-syn [27]. Our finding that the interaction between UBL3 and α-syn was downregulated by oxidative stress, prompted inquiries into its potential contribution to neurodegenerative processes. The downregulation of UBL3’s interaction with α-syn by oxidative stress promotes the α-syn free from the UBL3-α-syn complex disrupting the transfer of α-syn to the extracellular compartment by UBL3. This interferes with the potential role of UBL3 in preventing α-syn aggregation within the cell, leading to the accumulation of α-syn and the formation of aggregates.

In conclusion, our findings suggest that MGST3 plays a role in regulating the interaction between UBL3 and α-syn. Further investigation is needed to fully understand the underlying mechanisms involved in this process, and to determine how these factors may contribute to the development of neurodegenerative diseases.

## Reference

1. Downes, B.P.; Saracco, S.A.; Lee, S.S.; Crowell, D.N.; Vierstra, R.D. MUBs, a Family of Ubiquitin-Fold Proteins That Are Plasma Membrane-Anchored by Prenylation. J Biol Chem 2006, 281, 27145–27157, doi:10.1074/jbc.M602283200.

2. Ageta, H.; Ageta-Ishihara, N.; Hitachi, K.; Karayel, O.; Onouchi, T.; Yamaguchi, H.; Kahyo, T.; Hatanaka, K.; Ikegami, K.; Yoshioka, Y.; et al. UBL3 Modification Influences Protein Sorting to Small Extracellular Vesicles. Nat Commun 2018, 9, 3936, doi:10.1038/s41467-018-06197-y.

3. Raposo, G.; Stoorvogel, W. Extracellular Vesicles: Exosomes, Microvesicles, and Friends. J Cell Biol 2013, 200, 373–383, doi:10.1083/jcb.201211138.

4. Xia, X.; Wang, Y.; Zheng, J.C. Extracellular Vesicles, from the Pathogenesis to the Therapy of Neurodegenerative Diseases. Transl Neurodegener 2022, 11, 53, doi:10.1186/s40035-022-00330-0.

5. Goedert, M. Alpha-Synuclein and Neurodegenerative Diseases. Nat Rev Neurosci 2001, 2, 492–501, doi:10.1038/35081564.

6. Kawahata, I.; Finkelstein, D.I.; Fukunaga, K. Pathogenic Impact of α-Synuclein Phosphorylation and Its Kinases in α-Synucleinopathies. Int J Mol Sci 2022, 23, 6216, doi:10.3390/ijms23116216.

7. Goedert, M.; Jakes, R.; Spillantini, M.G. The Synucleinopathies: Twenty Years On. Journal of Parkinson’s Disease 2017, 7, S51–S69, doi:10.3233/JPD-179005.

8. Masliah, E.; Rockenstein, E.; Veinbergs, I.; Sagara, Y.; Mallory, M.; Hashimoto, M.; Mucke, L. β-Amyloid Peptides Enhance α-Synuclein Accumulation and Neuronal Deficits in a Transgenic Mouse Model Linking Alzheimer’s Disease and Parkinson’s Disease. Proceedings of the National Academy of Sciences 2001, 98, 12245–12250, doi:10.1073/pnas.211412398.

9. Klucken, J.; Shin, Y.; Masliah, E.; Hyman, B.T.; McLean, P.J. Hsp70 Reduces Alpha-Synuclein Aggregation and Toxicity. J Biol Chem 2004, 279, 25497–25502, doi:10.1074/jbc.M400255200.

10. Bin Chen; Mahmudul Hasan; Hengsen Zhang; Qing Zhai; A.S.M. Waliullah; Yashuang Ping; Chi Zhang; Soho Oyama; Mimi Mst. Afsana; Yuna Tomochika; et al. UBL3 Interacts with Alpha-Synuclein in Cells and the Interaction Is Downregulated by the EGFR Pathway Inhibitor Osimertinib. bioRxiv 2023, 2023.05.15.540732, doi:10.1101/2023.05.15.540732.

11. Jakobsson, P.-J.; Mancini, J.A.; Riendeau, D.; Ford-Hutchinson, A.W. Identification and Characterization of a Novel Microsomal Enzyme with Glutathione-Dependent Transferase and Peroxidase Activities*. Journal of Biological Chemistry 1997, 272, 22934–22939, doi:10.1074/jbc.272.36.22934.

12. Chen, J.; Xiao, S.; Deng, Y.; Du, X.; Yu, Z. Cloning of a Novel Glutathione S-Transferase 3 (GST3) Gene and Expressionanalysis in Pearl Oyster, Pinctada Martensii. Fish Shellfish Immunol 2011, 31, 823–830, doi:10.1016/j.fsi.2011.07.023.

13. Lu, L.; Pandey, A.K.; Houseal, M.T.; Mulligan, M.K. The Genetic Architecture of Murine Glutathione Transferases. PLoS One 2016, 11, e0148230, doi:10.1371/journal.pone.0148230.

14. Satoh, T.; Sakai, N.; Enokido, Y.; Uchiyama, Y.; Hatanaka, H. Free Radical-Independent Protection by Nerve Growth Factor and Bcl-2 of PC12 Cells from Hydrogen Peroxide-Triggered Apoptosis. J Biochem 1996, 120, 540–546, doi:10.1093/oxfordjournals.jbchem.a021447.

15. Jakobsson, P.J.; Morgenstern, R.; Mancini, J.; Ford-Hutchinson, A.; Persson, B. Membrane-Associated Proteins in Eicosanoid and Glutathione Metabolism (MAPEG). A Widespread Protein Superfamily. Am J Respir Crit Care Med 2000, 161, S20–24, doi:10.1164/ajrccm.161.supplement_1.ltta-5.

16. Bracalente, C.; Ibañez, I.L.; Berenstein, A.; Notcovich, C.; Cerda, M.B.; Klamt, F.; Chernomoretz, A.; Durán, H. Reprogramming Human A375 Amelanotic Melanoma Cells by Catalase Overexpression: Upregulation of Antioxidant Genes Correlates with Regression of Melanoma Malignancy and with Malignant Progression When Downregulated. Oncotarget 2016, 7, 41154–41171, doi:10.18632/oncotarget.9273.

17. Ayemele, A.G.; Tilahun, M.; Lingling, S.; Elsaadawy, S.A.; Guo, Z.; Zhao, G.; Xu, J.; Bu, D. Oxidative Stress in Dairy Cows: Insights into the Mechanistic Mode of Actions and Mitigating Strategies. Antioxidants (Basel) 2021, 10, 1918, doi:10.3390/antiox10121918.

18. Ashbrook, D.G.; Williams, R.W.; Lu, L.; Stein, J.L.; Hibar, D.P.; Nichols, T.E.; Medland, S.E.; Thompson, P.M.; Hager, R. Joint Genetic Analysis of Hippocampal Size in Mouse and Human Identifies a Novel Gene Linked to Neurodegenerative Disease. BMC Genomics 2014, 15, 850, doi:10.1186/1471-2164-15-850.

19. Castillo-Rangel, C.; Marin, G.; Hernández-Contreras, K.A.; Vichi-Ramírez, M.M.; Zarate-Calderon, C.; Torres-Pineda, O.; Diaz-Chiguer, D.L.; De la Mora González, D.; Gómez Apo, E.; Teco-Cortes, J.A.; et al. Neuroinflammation in Parkinson’s Disease: From Gene to Clinic: A Systematic Review. Int J Mol Sci 2023, 24, 5792, doi:10.3390/ijms24065792.

20. Lashuel, H.A.; Overk, C.R.; Oueslati, A.; Masliah, E. The Many Faces of α-Synuclein: From Structure and Toxicity to Therapeutic Target. Nat Rev Neurosci 2013, 14, 38–48, doi:10.1038/nrn3406.

21. Chen, B.; Hasan, Md.M.; Zhang, H. Interaction between UBL3 and Alpha-Synuclein in Cells Was Downregulated by the EGFR Pathway Inhibitor Osimertinib.

22. Hassan, W.; Noreen, H.; Rehman, S.; Kamal, M.A.; da Rocha, J.B.T. Association of Oxidative Stress with Neurological Disorders. Curr Neuropharmacol 2022, 20, 1046–1072, doi:10.2174/1570159X19666211111141246.

23. Barmaki, H.; Morovati, A.; Eydivandi, Z.; Jafari Naleshkenani, F.; Saedi, S.; Musavi, H.; Abbasi, M.; Hemmati-Dinarvand, M. The Association between Serum Oxidative Stress Indexes and Pathogenesis of Parkinson’s Disease in the Northwest of Iran. Iran J Public Health 2021, 50, 606–615, doi:10.18502/ijph.v50i3.5621.

24. Sian, J.; Dexter, D.T.; Lees, A.J.; Daniel, S.; Agid, Y.; Javoy-Agid, F.; Jenner, P.; Marsden, C.D. Alterations in Glutathione Levels in Parkinson’s Disease and Other Neurodegenerative Disorders Affecting Basal Ganglia. Ann Neurol 1994, 36, 348–355, doi:10.1002/ana.410360305.

25. Bellinger, F.P.; Bellinger, M.T.; Seale, L.A.; Takemoto, A.S.; Raman, A.V.; Miki, T.; Manning-Boğ, A.B.; Berry, M.J.; White, L.R.; Ross, G.W. Glutathione Peroxidase 4 Is Associated with Neuromelanin in Substantia Nigra and Dystrophic Axons in Putamen of Parkinson’s Brain. Mol Neurodegener 2011, 6, 8, doi:10.1186/1750-1326-6-8.

26. Sandeep; Sahu M.R.; Rani, L.; Kharat, A.S.; Mondal, A.C. Could Vitamins Have a Positive Impact on the Treatment of Parkinson’s Disease? Brain Sciences 2023, 13, 272, doi:10.3390/brainsci13020272.

27. Bosco, D.A.; Fowler, D.M.; Zhang, Q.; Nieva, J.; Powers, E.T.; Wentworth, P.; Lerner, R.A.; Kelly, J.W. Elevated Levels of Oxidized Cholesterol Metabolites in Lewy Body Disease Brains Accelerate α-Synuclein Fibrilization. Nat Chem Biol 2006, 2, 249–253, doi:10.1038/nchembio782.

