## Supplemental Table1 for "UBL3 interaction with α-synuclein is downregulated by silencing MGST3"

Supplemental Table 1

The list of siRNAs used in the current study

| **siRNA targeting** | **siRNA ID*** | **Sense 5′- 3′** | **Antisense 5′- 3′** |
| --- | --- | --- | --- |
| *QSOX1* | s224485 | CCU GCA CUC CGU GAA UGA Att | UUC AUU CAC GGA GUG CAG Gaa |
|  | s11505 | GGA UUC UUU GCG AGA AAU Att | UAU UUC UCG CAA AGA AUC Cat |
| *QSOX2* | s46761 | GAA UAU UCC UUA CUA AUC Att | UGA UUA GUA AGG AAU AUU Cca |
|  | s46763 | GAG GUG AUC UUA GAC CUG Att | UCA GGU CUA AGA UCA CCU Ccc |
| *HTATIP2* | s230128 | CGA CGA GGA AGC UUA UAA Att | UUU AUA AGC UUC CUC GUC Gaa |
|  | s230129 | AGA GUG CUC UUA AAG GAA Att | UUU CCU UUA AGA GCA CUC Ugc |
| *UBE3C* | s18661 | CAG GGU CUA UGG UAC CGU Utt | AAC GGU ACC AUA GAC CCU Gtg |
|  | s18660 | GAG ACA CUU UUG CGA AGU Att | UAC UUC GCA AAA GUG UCU Cgt |
| *COPS5* | s21627 | GCA AUC GGG UGG UAU CAU Att | UAU GAU ACC ACC CGA UUG Cat |
|  | s21629 | GGA UUG AUG UUA GUA CUC Att | UGA GUA CUA ACA UCA AUC Cca |
| MGST3 | s8762 | AGA ACA CGU UGG AAG UGU Att | UAC ACU UCC AAC GUG UUC Ugg |
|  | s8764 | GAC GAG UUC UUU AUG CUU Att | UAA GCA UAA AGA ACU CGU Cca |
| *NSF* | s181 | GUU ACA UUA UGA ACG GUA Utt | AUA CCG UUC AUA AUG UAA Ctt |
|  | s182 | GGC AGA CUU UCU ACA UGC Att | UGC AUG UAG AAA GUC UGC Ctt |
| *HECTD1* | s24575 | GGC UCG AGA UUU AUA CGA Utt | AUC GUA UAA AUC UCG AGC Cat |
|  | s24576 | CAG UUG CGU UAA UUC GAA Att | UUU CGA AUU AAC GCA ACU Gct |
| *SAE1* | s19551 | GUU CCG UAC AGA UAA AGG Att | UCC UUU AUC UGU ACG GAA Ctt |
|  | s19552 | GCU GGA UCA CGA ACA GGU Att | UAC CUG UUC GUG AUC CAG Cat |
| *ATG3* | s34731 | GAA UAA CGG AAG CCG UUA Att | UUA ACG GCU UCC GUU AUU Cct |
|  | s34733 | GCG GAU GGG UAG AUA CAU Att | UAU GUA UCU ACC CAU CCG Cca |

*Silencer Select (Ambion, Life Technologies)
